# Nonlinear Parameter and State Estimation Approach in End-stage Kidney Disease Patients

**DOI:** 10.1101/2022.04.02.486844

**Authors:** Rammah M. Abohtyra, Tyrone L. Vincent

**Affiliations:** Department of Engineering Science and Mechanics, The Pennsylvania State University, University Park, State College, 16802, Pennsylvania, USA; Department of Electrical Engineering, Colorado School of Mines, 1500 Illinois St, Golden, 80401, Colorado, USA

## Abstract

**Background:** Blood and fluid volume management in End-stage Kidney Disease (ESKD) patients plays an essential role in dialysis therapy to replace kidney function. Reliable knowledge of blood and fluid volumes before and during dialysis could be used to improve treatment outcomes significantly.

**Objective:** This study aims to develop an estimation approach providing predictable information on blood and fluid volumes before and during a regular dialysis routine.

**Methods:** A new approach is developed to estimate blood volume, fluid overload, and vascular refilling parameters from dialysis data. The method utilizes a nonlinear fluid volume model, an optimization technique, and the Unscented Kalman Filter (UKF) incorporated with data. This method does not rely on restricted ultrafiltration (UF) and dilution protocols and uses the Fisher information matrix to quantify error estimation.

**Results:** Accurate estimations for blood volumes (5.9±0.07L and 4.8±0.03L) and interstitial fluid volumes (18.81±0.15L and 12.19±0.03) were calculated from dialysis data consisting of constant and stepwise UF profiles. We demonstrated that by implementing the estimated parameters into the model, a precise prediction of the measured hematocrit (HCT) can be achieved during the treatment.

**Conclusion:** We showed that the result does not depend highly on initial conditions and can be accurately estimated from a short data segment. A new method, applicable to the current dialysis routine, is now available for ESKD patients to be implemented within the dialysis machines.

## 1 Introduction

End-stage kidney disease (ESKD) is a major human health concern, leading to a significant deficit in the quality of life, higher mortality, and substantial economic impact [1]. ESKD patients receive 3-4 dialysis sessions (each session is 2-5hrs long) per week. During dialysis, ultrafiltration (UF) removes excess fluid to reach dry weight [2, 3]. Intradialytic hypotension (IDH) is a common adverse outcome and is associated with a higher risk of cardiovascular morbidity and rate of death [4–7]. Therefore, knowing a patient’s blood and fluid volumes at the start of dialysis and during UF could allow clinicians to better guide fluid balance and return ESKD patients to their dry weight safely [8, 9].

This work aims to develop a new estimation approach to estimate, from dialysis data, blood and fluid volumes before dialysis and predict the change in these volumes during UF. The method also estimates critical fluid refilling-related parameters and predicts the patient’s hematocrit (HCT) precisely when these parameters are estimated correctly. One primary goal of this method is to detect IDH by monitoring the blood volume change during UF. We note that our approach does not rely on the currently restricted UF rate protocols (such as pulse-UF rate, decrease/increase-UF rates, etc.) for estimating fluid overload and an exogenous bolus infusion of 240 mL dialysate injected into the extracorporeal blood circulation to estimate blood volume. Instead, the method utilizes dialysis data generated in a regular dialysis routine using standard UF profiles, including constant and stepwise profiles.

The motivation for this work is fourfold. Firstly, overcome the limitation that the current estimation of blood volume and fluid overload depends on restricted protocols of UF rates and dialysate dilutions. Secondly, accurate estimation of blood volume and fluid overload at the start of dialysis would assist in determining an appropriate dry weight. Thirdly, personalized predictions for the changes in blood and fluid volumes during UF could help to prevent IDH, assess fluid refilling rate, and ensure the achievement of dry weight. Lastly, providing a new technique to be integrated within the current dialysis machines to provide reliable measurements for blood and extracellular fluid volumes, which cannot be provided clinically. We note that since Hemodialysis (HD) is one type of dialysis, this method is applicable for HD patients.

Predictable knowledge of blood volume during UF could substantially improve blood volume management and reduce IDH. In ESKD patients, blood volume is essential in the long run because it has been shown that extra blood volume mediates most of the cardiovascular damage, including vascular stiffening, left ventricular hypertrophy, and congestive heart failure [8]. On the other hand, a decline in blood volume primarily affects the venous side of the systematic circulation, alters the cardiac filling pressure, impacts cardiac output, and then affects arterial blood pressure [6]. During dialysis treatment therefore the progression of IDH is mainly due to a reduction in blood volume when the UF rate exceeds the vascular refilling rate, the rate of fluid transfer from the interstitial compartment into the intravascular space [10]. Without UF, the appearance of IDH is not common, indicating that UF is the leading cause of IDH [8].

Over decades, noninvasive methods have been developed to estimate blood volume using the relative blood volume changes (RBV) [11]. Later, the infusion dialysate technique was developed as a new technique to estimate blood volume during a dialysis routine [2, 12–14]. However, all these methods require an indicator dilution (fluid bolus, 240 mL) of dialysate infused to a patient over a short time to estimate the blood volume at the time of the bolus infusion. Therefore, the estimated blood volume does not precisely correspond to the blood volume before dialysis [13]. Other methods based on compartment mathematical models have been developed to estimate blood volume using measurements of RBV and controlled UF protocols [8, 12, 13]. The concern with these methods is that these methods are based on restricted UF protocols, which are difficult to apply in every normal dialysis routine, and a systemic hemoglobin measurement that must be converted to a whole-body hemoglobin concentration to obtain an accurate blood volume estimate [8].

Dry weight and fluid overload are used to explain body fluid status [15]. The concept of dry weight in an ESKD patient is defined as the patient without excess fluid volume at post-dialysis weight [16]. On the other hand, fluid overload can be generally defined as the extracellular fluid volume (EFV) accumulated in the body of greater than a normal range [17]. But the normal range of EFV is uncertain. This uncertainty in the normal range of EFV, the variability of excess fluid in ESKD patients due to age and gender, kidney disease progression, and the deficiency of reliable techniques to measure EFV made the assessment of fluid overload a challenging task [18].

The current methods for measuring fluid overload can be generally divided into three techniques: static, dynamic, and a combination of both methods [19]. In the static methods, healthy subjects are used as references to evaluate fluid overload [19–21]. However, these techniques cannot provide a quantitative value of fluid overload for ESKD patients [19]. Two methods are mainly used for the dynamic methods: the RBV-based method and calf bioimpedance spectroscopy (cBIS) [19]. In the RBV method, a measurement displays a change in plasma volume, but it cannot provide direct information about fluid overload in the interstitial compartment. The main advantage of the cBIS method is that it monitors the EFV changes, including plasma and interstitial fluid volume, and does not require a reference parameter to estimate fluid overload. However, this method could be affected by lower limb venous thromboses [19]. The combination of static and dynamic methods can provide reliable information on the dry weight estimation of an ESKD patient. However, these methods may take a long time for the post-dialysis weight to be reduced slowly when the patient’s EFV is overloaded [19].

The vascular refilling rate during UF varies throughout the dialysis session, affects the blood volume drop, and relies on Starling’s driving forces (hydrostatic and osmatic pressure gradients across the capillary wall) [22]. These driving forces can be influenced by arteriolar tone, venous tone, changes in plasma solute concentrations (i.e., acid-base, electrolytes, and proteins), capillary permeability, and the overall patient’s fluid volume state [22]. The change in the vascular refilling rate is based on the degree of interstitial fluid overload, so this rate is, on average higher in an ESKD patient with a higher degree of fluid overload and lower in patients whose fluid status is closer to their dry weight. As a result, blood volume can vary widely between ESKD patients and even within the same patient over time [8, 23].

It is not possible to have direct measures for the vascular refilling rate during dialysis treatment [24, 25]. However, an estimation for the vascular refilling rate during dialysis can be used to obtain an adequate balance between UF and interstitial fluid [6]. Therefore, model-based methods have been developed to estimate the vascular refilling rate using the RBV trajectory and are based on the assumption that the vascular refilling rate depends only on the capillary membrane and UF [8, 25–28]. The problem is that the UF used in these methods is guided by restricted UF protocols, affecting the achievement of dialysis goals [28]. Therefore, new techniques proper to the standard UF are suggested.

To this end, we develop a new method to overcome the above limitations. This method uses dialysis data to estimate vascular refilling-related parameters, blood volume, and extracellular fluid overload before dialysis. We depend on the UKF and the model to predict the changes in these volumes in response to UF and detect IDH events. The method is not reliant on the current restricted UF and dialysate infusion protocols.

The paper is organized as follows. Section II summarizes the nonlinear model. Section III introduces a new estimation method, including parameter estimation algorithm, prediction algorithm, and estimation sensitivity analysis. Section IV illustrates the results using our method with clinical data. Section V provides discussion and future directions, and the conclusions are made in Section VI.

## 2 Materials and Methods

The proposed approach is derived in this section. This approach combines a nonlinear fluid volume model, an optimization method with an objective function, and the UKF incorporated with data. In what follows, we will introduce these components.

### 2.1 Two Compartment Fluid Volume Model

The model is given by a validated two-compartment fluid volume model defined by nonlinear continuous-time differential equations and used for ESKD patients. Previously, we have successfully used the linearized version of this model to develop a personalized UF method [10, 29, 30], but here we use the ‘full’ nonlinear model.

We provide a brief summary about the nonlinear two-compartment model illustrated in Fig. 1. The model captures the EFV dynamics, which comprises of the intravascular (plasma) and interstitial compartments, separated by a capillary membrane. The model includes the lymphatic flow, which returns a portion of fluid from the interstitial space into the intravascular space. The states of the model are the plasma volume, *V*_*p*_ (L), and the interstitial fluid volume, *V*_*i*_ (L); while the input, *u*, is the UF rates (UFRs) (L/min). The output is the HCT, which is the proportion of the total blood volume (*V*_*b*_ = *V*_*rbc*_ + *V*_*p*_) that consists of red blood cells (RBCs) volume, *V*_*rbc*_ (L), i.e., *HCT* = *V*_*rbc*_*/V*_*b*_, which is usually expressed as a percentage in clinical practices but is expressed here as a fraction.

**Fig. 1.**
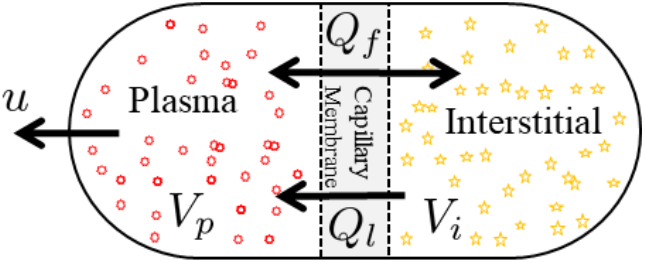
The two-compartment model consists of interstitial fluid volume (*V*_*i*_) and intravascular (plasma) fluid volume (*V*_*p*_) separated by a capillary membrane; microvascular flow (*Q*_*f*_), lymph flow (*Q*_*l*_), and UF (*u*).

The microvascular flow, *Q*_*f*_ (refilling/filtration), describes the fluid exchange between the intravascular and interstitial spaces through the capillary membrane. The mathematical description of this model during dialysis is given by the nonlinear differential system:

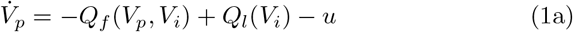

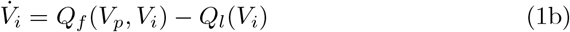

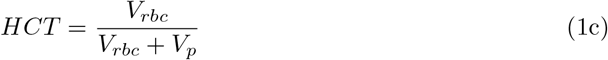

where *Q*_*f*_ (L/min) represents the microvascular (shift/refill) filtration flow, in which, *K*_*f*_ (L/min.mmHg) is the transcapillary permeability coefficient; and *Q*_*l*_ (L/min) represents the lymphatic flow:

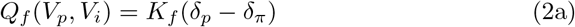

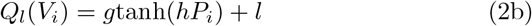

where *d*_*p*_ and *d*_*π*_ represent the hydrostatic and osmotic pressure gradients, respectively:

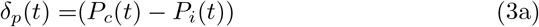

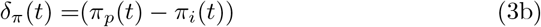

where *P*_*c*_, *P*_*i*_, *π*_*p*_, *π*_*i*_ refer to the hydrostatic capillary pressure, interstitial pressure, plasma colloid osmotic pressure, and interstitial colloid osmotic pressure, respectively; and *g, h*, and *l* are constants. The equations of these components, nominal parameters, and constants are provided in the Appendix.

### 2.2 Patient Fluid Volume Variability

In ESKD patients, the variability in blood and fluid volumes is high and depends on several factors, including fluid compartment sizes, fluid overload, fluid shift/refill between interstitial and intravascular spaces [22], and patient’s characteristics. Here, we identify six parameters of the model (1) as patient-specific parameters to be estimated from patient’s data.

To estimate fluid overload, the initial states of the model are estimated, including plasma and interstitial fluid volumes *V*_*p*_(0), *V*_*i*_(0), respectively. Additionally, the initial (absolute) blood volume *V*_*b*_(0) can be determined by estimating first *V*_*rbc*_(0) and then adding *V*_*b*_(0) = *V*_*p*_(0)+*V*_*rbc*_(0). Since UF does not remove red blood cells, *V*_*rbc*_ will be fixed during dialysis [31]. Therefore, the change in *V*_*p*_ influences blood volume.

During UF, the capillary filtration coefficient *K*_*f*_ regulates the fluid refilling rate. In contrast, the hydrostatic capillary pressure parameters (*d*_1_ and *d*_2_) promote fluid movement from the intravascular space to interstitial space and control the fluid shift/refill across the capillary membrane wall. These parameters vary among ESKD patients [22, 30].

Since the lymphatic system’s parameters are insensitive to HCT and the system returns a small amount of fluid to the intravascular space, these parameters are fixed for all patients. Thus, the patient-specific parameters are included in this vector: *θ* = [*V*_*p*_(0), *V*_*i*_(0), *V*_*rbc*_(0), *K*_*f*_, *d*_1_, *d*_2_]^*T*^, whereas, the other parameters in Table A1 are nominal for all patients.

### 2.3 Objective Function and Data

The objective function (4) is defined to optimize the parameters *θ* by minimizing the sum of the square errors between HCT measurements (*z*_*i*_) and the model’s HCT (*y*(*θ, t*_*i*_)) (1) in response to the UF as an input to the model, where *t*_*i*_ indicates the time of HCT measurements (*i* = 1, …, *N*). The objective function is given by

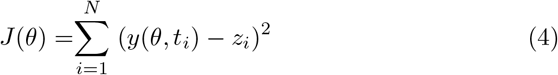

To test the method, two dialysis data sets consisting of HCT measurements taken from the Crit-Line Monitor and two UF profiles, including constant and stepwise UF profiles, widely used in dialysis practices [32, 33], are used. The data were published in this previous study [34].

In this data, the rate in the constant UF profile is fixed at 600 mL/hrs, with a dialysis goal to remove 2 L within 203 min. In the stepwise profile, the UF rate is started at 750 mL/hrs, then increased from 880 to 1160 mL/hrs, and then decreased from 1080 to 940 mL/hrs, with a dialysis goal to remove 3.86 L within 234 min.

## 3 New Estimation Approach

The main contribution of this paper is the development of a new approach for ESKD patients to estimate the physiological parameters 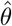. The uncertainty in the estimation is studied based on the Fisher information matrix. Further, the sensitivity of the estimation to an initial guess is explored using random initial conditions.

The estimation approach consists of parameter and state estimation algorithms performed separately in sequence (see Fig. 2).

**Fig. 2.**
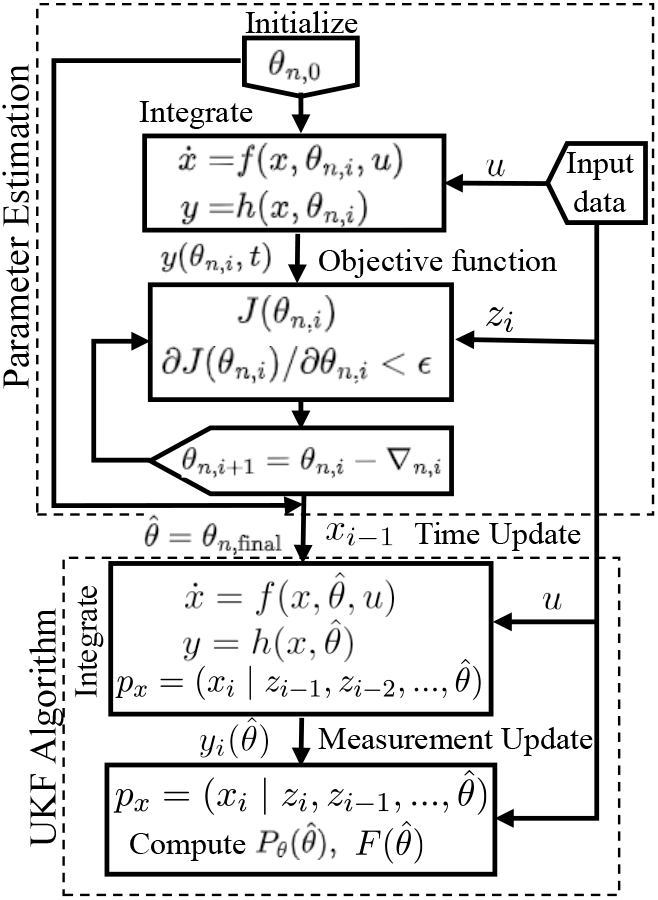
A schematic representation for Algorithm 1. The parameter estimation and prediction (UKF) Algorithms are independently performed. Note that the estimated parameter 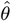 is an input to the UKF Algorithm.

### 3.1 Parameter Estimation Algorithm

The estimation approach utilizes a general nonlinear model of continuous-time differential equation described by a state transition (*f*) and observation function (*h*), given by these equations:

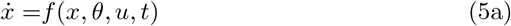

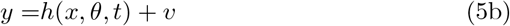

where *x* ∈ ℝ^*n*^ denotes the state of the system; *u* denotes the input to the system; and *θ* ∈ ℝ^*p*^ contains the model parameters whose values (or some of them) are estimated from data; *y* is the model output; *t* is the time *t*_0_ *≤ t ≤ t*_*f*_ between initial, *t*_0_, and final time, *t*_*f*_ ; and *v* represents measurement noise.

The algorithm minimizes the objective function (4) using the least squares to compute an optimal solution 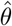

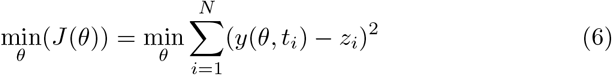

where *z*_*i*_ and *y*(*θ, t*_*i*_) are defined in (4) (refer to observed measurements and output trajectory of the nonlinear model). The optimal solution of (6) is denoted by 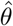.

The proposed estimation algorithm, as shown in Fig. 2, consists of two nested loops: the outer one loops over a random set of initial conditions *θ*_*n*,0_, and the inner loop is based on the Levenberg-Marquardt method where the value *θ*_*n*_ is updated on the *i*th cycle by *θ*_*n,i*+1_ = *θ*_*n,i*_ *−* ∇_*n,i*_ where ∇_*n,i*_ uses the steepest descent method [35]. The MATLAB function ‘lsqcurvefit’ is used to implement the inner loop.

Note that, as described in Fig. 2, the estimated parameter 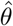 is an input to the UKF Algorithm, which is demonstrated in the next subsection (3.2).

### 3.2 Prediction Algorithm

The UKF [36] is integrated with the nonlinear model and data to compute the estimation error using the Fisher information matrix, update states (blood and fluid volumes), and predict the measured HCT during UF, as detailed in this section.

In order to estimate parameter uncertainty and make predictions of state trajectories, using the UKF, we introduce model uncertainty (*n*) into (5a) to come

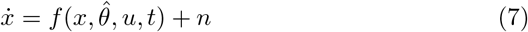

and made the assumption that the distribution of the model uncertainty (*n*) and measurement noise (*v*) in (5b) is piece-wise constant over intervals of length *T*_*s*_ seconds (*T*_*s*_ is a sampling time), with the magnitude for interval *k* (*k* is an index) a Gaussian random vector with zero mean given by

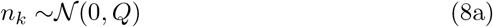

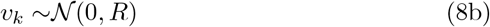

where 𝒩 represents the notation for Normal Gaussian Distribution; *Q* and *R* are the covariance of model uncertainty and measurement noise, respectively. Consequently, we assume *x* is Gaussian. Since *y* is conditionally affected by the state *x*, we also assume that *y* is Gaussian. Thus, the conditional state and output density functions are Gaussian:

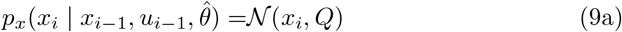

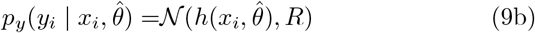

where *x*_*i*_ is the solution to (5) with *n* = 0, initial condition *x*_*i−*1_ and input *u*_*i−*1_.

As illustrated in Fig. 2, the prediction algorithm uses the UKF with the estimated parameter 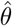 and the nonlinear model (5) to compute the Fisher information matrix 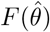, update states, and predict the output of this model. Here, we provide iterative steps for updating the UKF, utilizing observations *z*_0_, *z*_1_, …, *z*_*N*_ at a sampling time *T*_*s*_, and an input *u*, where *u* is held as a piecewise constant during the interval (*i −* 1)*T*_*s*_ *≤ t ≤ iT*_*s*_:

- 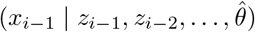 represents the previous estimated state.
- 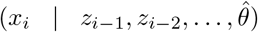 represents the prediction update step (time update), obtained by integrating model (5) between time *t* = (*i −* 1)*T*_*s*_ to *t* = *iT*_*s*_, with initial conditions 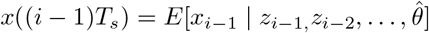.
- 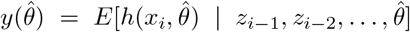 represents the prediction step of a measurement *z*_*i*_.
- 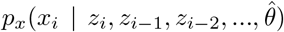 represents the measurement update step of the UKF.

Algorithm 1 below presents the parameter estimation (A) and prediction (UKF) algorithm (B).

### 3.3 Parameter Estimation Sensitivity Analysis

We use the Fisher information matrix [37] to quantify the quality of the estimation due to the model uncertainty and measurement noise. This analysis uses the following assumptions.

- The distribution of *y* can be approximated by a Gaussian distribution.
- The experimental uncertainty is captured by model (5)
- The covariance of the prediction error is independent of *θ*.
- The UKF is approximately optimal, so that the prediction errors are uncorrelated.
- The parameter estimates are unbiased

#### Algorithm 1: The Parameter and prediction Algorithms

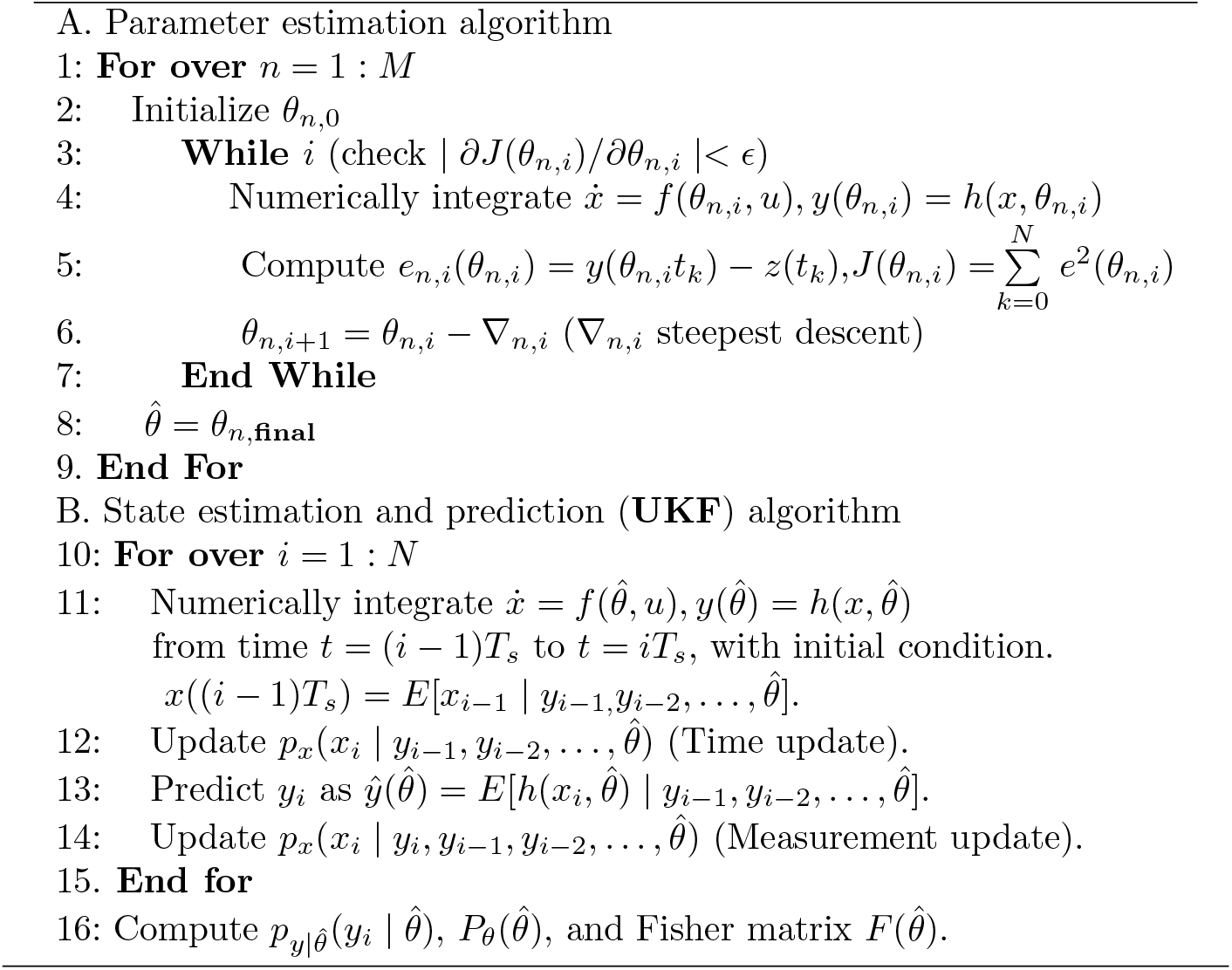

#### Theorem 1

*Under the assumptions listed above, the covariance of the parameter estimates (P*_*θ*_*) is bounded below by*

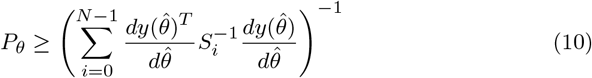

*where S*_*i*_ *is the covariance of the prediction error calculated using the UKF*.

**Proof:** For unbiased parameter estimates, the parameter estimate covariance is bounded below by the inverse of the Fisher information matrix (see e.g. [37]), which is calculated from the likelihood function. The likelihood function for the measurements *Z* where *Z* = [*z*_0_ ⋯ *z*_*N*_]^*T*^ is given by

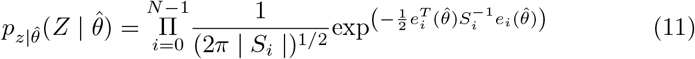

where 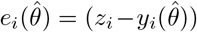, in which 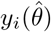 is the UKF one step ahead prediction; and *S*_*i*_ = *HP*_*i*_*H*^*T*^ + *R* is the covariance of the error 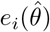, where *P*_*i*_ is the estimated state error covariance at time *i*, and *H* is the mapping from state to output (i.e., *H* = [1 0] for the model (1)). Then the log-likelihood function is

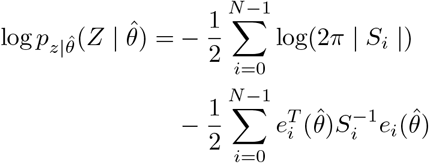

Then

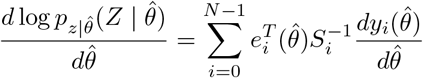

The Fisher information matrix is given by the covariance of this expression, so we find

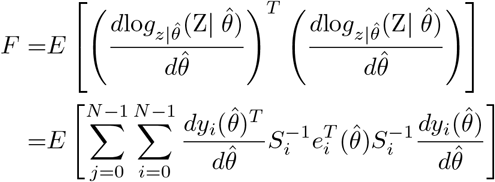

Under the assumption that the prediction error 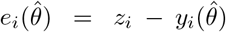 is uncorrelated,

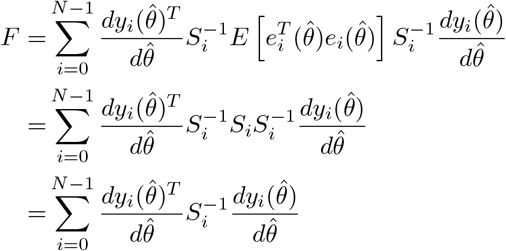

The result follows by inverting the Fisher information matrix. □

### 3.4 Parameter Initialization Sensitivity

To address the sensitivity to initial conditions (initial guess for *θ*), we investigate the uncertainty and uniqueness of the optimal solution, 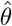, using randomly sampling a set of initial parameters Θ_0_ (uniform distribution) within bounded intervals. We choose the boundaries of the intervals to exclude unrealistic parameter variations and include all possible solutions.

For model (1), we compute the upper and lower boundaries of the intervals of the parameters (*V*_*p*_(0), *V*_*i*_(0), *V*_*rbc*_(0), *K*_*f*_, *d*_1_, *d*_2_) within plausible physiological ranges centered around the nominal values of the parameters in

Table A1, using ±50% of the nominal values, which yields the set of vectors Θ_0_ = [4.5, 1.5] × [16.5, 5.5] × [3, 1] × [0.0086, 0.0029] × [0.015, 0.005] × [2.19, 0.73]. In this set, we use the nominal volumes, *V*_*p*_ = 3, *V*_*i*_ = 11, and *V*_*rbc*_ = 2 (L), used in [29].

The output of this computational method is a distribution of optimal solutions represented by a probability density, which allows to understand how informative, unique, and uncertain a given estimation is. Therefore, if the distribution of the solutions is narrow or wide, we can conclude that the estimation either does or does not depend highly on the given initial parameters Θ_0_.

## 4 Results

In this section, the method was applied to clinically measured dialysis data, including HCT measurements and UF (constant and stepwise) profiles described in Section (2.3) to test the performance of the method.

### 4.1 Application to Dialysis Data

#### 4.1.1 Constant UF Profile

The constant UF profile and HCT measurements were shown in Fig. 3 (A blue, B black). This profile was applied to remove 2 L of fluid, within 203 min, and at the rate of 600 mL/hrs. The nominal values *θ*_0_ = [3, 11, 2, 0.0057, 0.01, 1.46]^*T*^ was used as an initial guess for estimation.

**Fig. 3.**
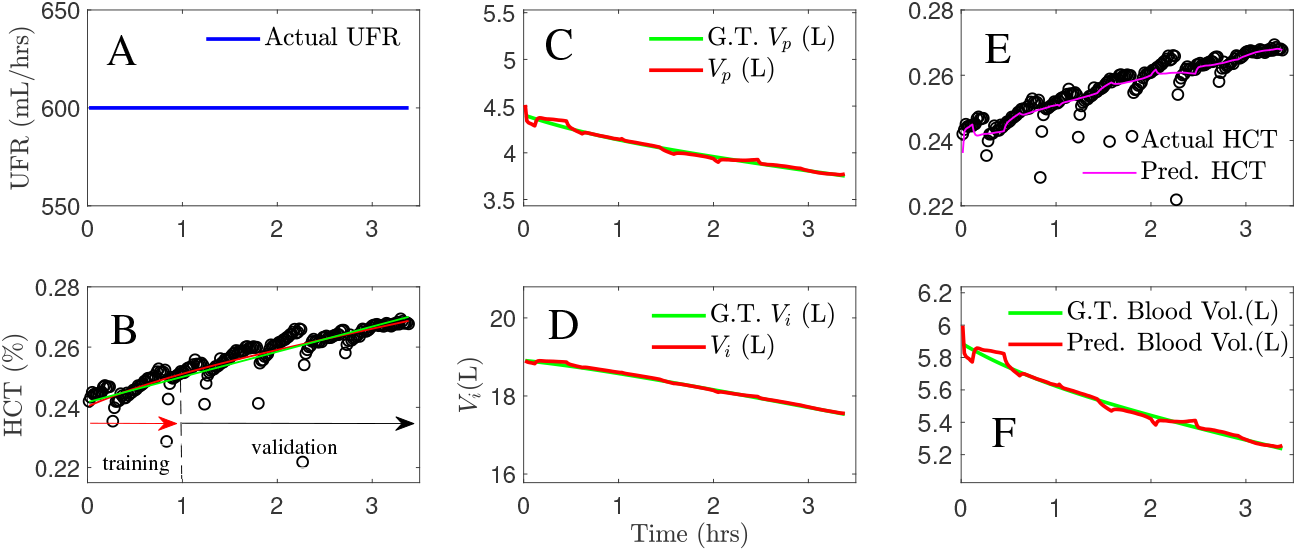
Estimation details using the constant UF rate (UFR) profile (A, blue) and measured HCT (B, black), model HCT (B, red) generated from all data points and model HCT (B, green) generated using the first (0-60 min) data segment for training and the rest for validation. Personalized predictions for the measured HCT using the UKF algorithm (E, magenta), and fluid and blood volume updates (C, D, F) for plasma volume (C, red), interstitial volume (D, red), and blood volume (F, red) versus ground truth (GT) trajectories (green).

The first (0-60 min) data segment was utilized to train the model, estimate the parameters, and validate the rest of the data. Table 1 shows the results *θ*_2_. In addition, all data points were used for estimation 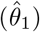. Fig. 3 B shows a comparison between the modeled HCTs (red) with 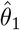 and (green) with 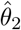, under the constant UF profile. A good agreement was observed between the measured HCT (B, black) and the modeled HCTs (red, green).

**Table 1.** Estimation results were computed using dialysis data, for all data points (*θ*_1_) and compared with the first data segment 0-60 min and 0-90 min (*θ*_2_) using constant and stepwise UF profiles, respectively.

| Parameters | (all data set)<br>$\hat{\theta}_1 \pm \text{SD}$ | 95% Confidence interval (CI) | diag($F^{-1}$) | (0-60, 90)min<br>$\hat{\theta}_2 \pm \text{SD}$ |
| --- | --- | --- | --- | --- |
| Example 1 (Standard UFR) |  |  |  |  |
| $V_p(0)$ | $4.46 \pm 5\text{e-}3$ | [4.45, 4.47] | $2.7\text{e-}5$ | $4.47 \pm 5.5\text{e-}3$ |
| $V_i(0)$ | $18.81 \pm 0.15$ | [18.5, 19.1] | 0.02 | $18.82 \pm 0.23$ |
| $V_{rbc}$ | $1.42 \pm 0.06$ | [1.31, 1.54] | $3.1\text{e-}3$ | $1.43 \pm 0.08$ |
| $K_f$ | $1.5\text{e-}3 \pm 4\text{e-}4$ | [0.7e-3, 2.4e-3] | $1.7\text{e-}7$ | $1.5\text{e-}3 \pm 6\text{e-}4$ |
| $d_1$ | $1.36 \pm 0.52$ | [0.34, 2.37] | 0.27 | $1.36 \pm 0.73$ |
| $d_2$ | $0.02 \pm 0.06$ | [-0.09, 0.13] | $3.3\text{e-}3$ | $0.02 \pm 0.08$ |
| Example 2 (stepwise UFR) |  |  |  |  |
| $V_p(0)$ | $3.27 \pm 5\text{e-}4$ | [3.26, 3.28] | $2.1\text{e-}5$ | $3.27 \pm 5.2\text{e-}3$ |
| $V_i(0)$ | $12.19 \pm 0.03$ | [12.13, 12.25] | $7.78\text{e-}4$ | $12.19 \pm 0.12$ |
| $V_{rbc}$ | $1.49 \pm 0.03$ | [1.44, 1.56] | $8.7\text{e-}4$ | $1.49 \pm 0.1$ |
| $K_f$ | $2.2\text{e-}3 \pm 5\text{e-}4$ | [1.3e-3, 3.2e-3] | $2.3\text{e-}7$ | $2.2\text{e-}3 \pm 1.3\text{e-}3$ |
| $d_1$ | $1.06 \pm 0.36$ | [0.36, 1.77] | 0.13 | $1.1 \pm 0.94$ |
| $d_2$ | $7.9\text{e-}3 \pm 0.02$ | [-0.029, 0.045] | $3.5\text{e-}4$ | $7.9\text{e-}3 \pm 0.048$ |

The uncertainty around each estimated parameter was defined by 95% CIs as shown in Table 1. The estimated variances were computed from diag 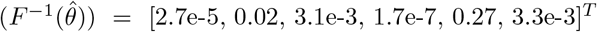 (Theorem 1). In Table 1, narrow CIs were obtained for *V*_*p*_(0), *V*_*i*_(0), *V*_*rbc*_, and *K*_*f*_, ensuring the capability of the approach in estimating these parameters precisely. However, the CIs for *d*_1_ and *d*_2_ were widely spread due to the insensitivity of these parameters to HCT measurements.

The distribution of 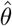 was computed using the initial set Θ_0_ defined in Section (3.4). Fig .4 shows the distribution using the Kernel Probability Density (KPD) calculated by the function ‘ksdensity’ in MATLAB. Therefore, the KPD showed that the solution is virtually normally distributed around the set 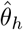 which has the highest density (peak), 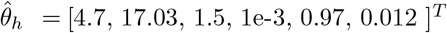.

**Fig. 4.**
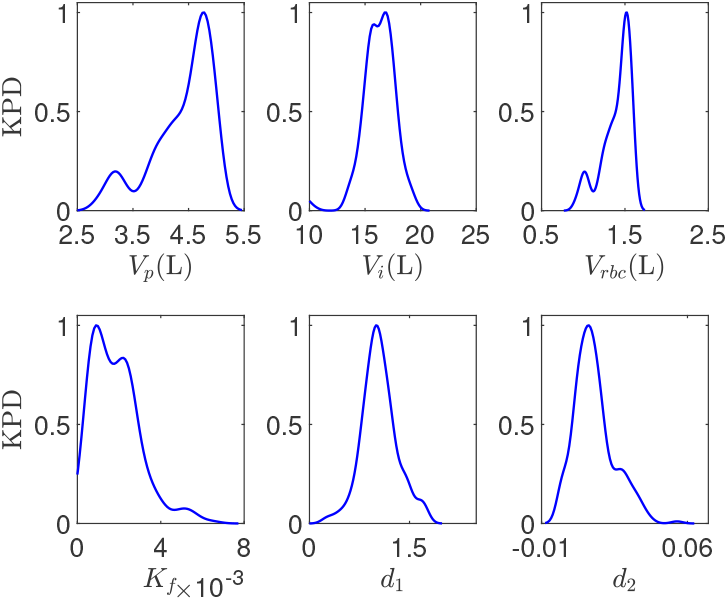
The Kernel Probability Density (KPD) estimate for the estimated parameters, under the constant UF profile, over the random initial solutions Θ_0_. The highest density parameter set is 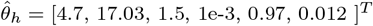.

Figure 3 shows the performance of the prediction algorithm under the constant UF profile, in which, when the estimated physiological parameters (*K*_*f*_, *V*_*rbc*_,*d*_1_,*d*_2_) were implemented in the model, a precise prediction for the measured HCT (E, magenta) was obtained. In addition, updates for the blood and fluid volumes (C, D, F, red) were computed and compared with ground truth (GT, green) trajectories. These trajectories were generated from the model with the estimated parameters 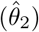 to represent the known patient’s true blood and fluid volumes.

The results in Table 1 indicated that the patient under the constant UF was in an overload status, which can be understood by the initial volumes of plasma (4.46±5e-3 L), interstitial fluid (18.81±0.15 L), and blood (5.9±0.07L), compared with these studies [14, 23, 38]. However, the vascular refilling rate was large, confirmed by the large refilling filtration coefficient (1.5±e-3) [39], indicating that during the constant UF, a large amount of interstitial fluid was shifted to the plasma space. Furthermore, the estimated volume of the red blood cells (1.42±0.06) was low compared with this study [40].

#### 4.1.2 Stepwise UF Profile

The stepwise UF profile shown in Fig. 5 (A) was to remove 3.86 L of fluid within 234 min.

**Fig. 5.**
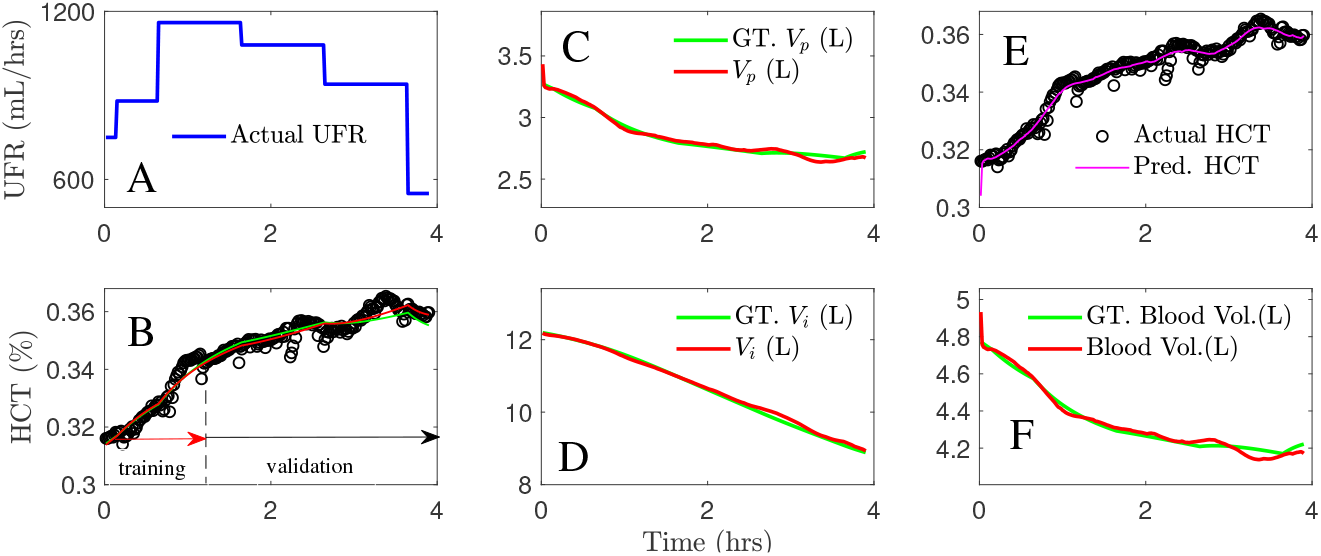
Estimation details for the stepwise UF rates (UFR) profile (A, blue) and measured HCT (B, black), and simulated HCT (B, red) from all data points and simulated HCT (B, green) using (0-90min) for training and the rest for validation. Personalized predictions for the actual HCT using the UKF (E, magenta), and volume updates (C, D, F) for plasma volume (C, red), interstitial volume (D, red), and blood volume (F, red) versus ground truth (GT) trajectories (green).

A data segment (0-90 min) was utilized to capture the dynamics and estimate the parameters 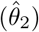. In addition, all data points were utilized to estimate the parameters 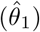 (see Table 1). The difference between the measured HCT and the modeled HCTs using 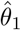 and 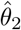 was negligible. Table 1 depicts the estimated variances, diag 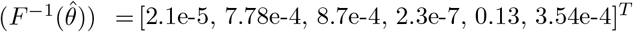.

Similar to the previous result, Fig. 6 shows that the KPD of 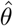 estimated over the initial set Θ_0_ was normally distributed around the parameter set that has the highest density, 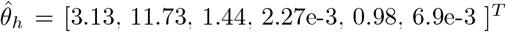. This parameter set was very close to 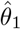 and 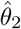 estimated using the nominal values.

**Fig. 6.**
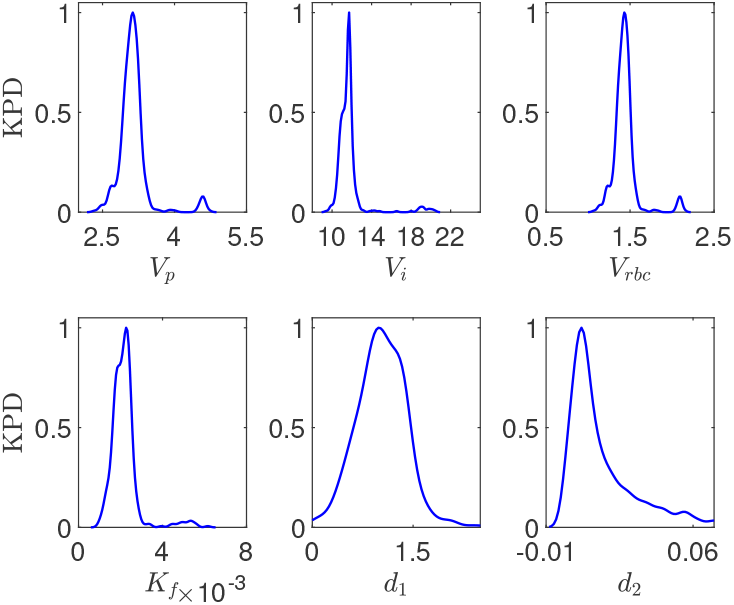
The Kernel Probability Density (KPD) estimate for the estimated parameters, under the stepwise UF, over the random initial solutions Θ_0_. The highest density parameter set is 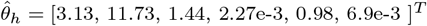.

A personalized prediction for the measured HCT (Fig. 5 E, magenta) was achieved even though the stepwise UF profile applied different removal rates during this treatment. Moreover, the updated fluid and blood volumes were shown in Fig. 5 (C, D, F, red) and compared with GT trajectories (green) generated from the model (1) with 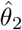 under the stepwise UF profile.

The estimation results in Table 1 showed that the extracellular fluid (plasma, interstitial) volume was overloaded (3.27±5e-4 L) and (12.19±0.03 L). However, the initial blood volume was in the normal range (4.8±0.03L) [14]. At the end of dialysis, the interstitial and plasma volumes were reduced from 12.19L to 8.9L and 3.27L to 2.8L, respectively (see Fig. 5 C, D). This observation confirmed that the largely removed fluid overload was driven from the interstitial fluid [8].

## 5 Discussion and Future Directions

The study presents a new method to estimate blood and fluid volumes and vascular refilling-related parameters from dialysis data. This study shows that the blood volume and extracellular fluid overload before starting dialysis can be successfully estimated from a small part of the data. When the estimated parameters are implemented into the model, predictions for these volumes and the measured HCT can be generated during UF. Our method is applicable to the standard UF profiles and independent of the dilution infusion protocols.

We note that dialysis has other purposes, including a correction for acidbase derangements, which utilizes different types of physiological mathematical models [41, 42]. This is beyond the scope of this study.

### 5.1 Validation

The accuracy of the estimation is validated using the Fisher information matrix, demonstrated in Theorem 1, in which the variance was calculated by taking the square root of the diagonal terms of the matrix. We showed that the initial blood and extracellular fluid volumes and the refilling coefficient can be estimated within a narrow range of uncertainty. However, due to the insensitivity of the hydrostatic capillary parameters to the HCT measurements, the CIs of these parameters were wide compared with the other parameters. We also evaluated the method using a data segment for training the model and estimating the parameters and then the rest of the data for validating the estimation. Further, another test compares the estimation using all data points with the estimation using the initial data segment. We showed that the difference between the estimated model’s responses is tiny.

Further, the fidelity of our prediction algorithm was tested by predating the measured HCT using the model’s HCT. Therefore, when the estimated parameters were implemented in the model, precise predictions for the measured HCT, under the UF profiles, were achieved even though the stepwise UF profile applied different removal rates during the treatment.

We analyzed the sensitivity of the estimation result to initial conditions to generate a probability distribution. The distribution of the estimated solutions, under both UF profiles, was approximately narrow and normally distributed around the solution with the highest density. Therefore, we concluded that the estimation result does not depend highly on initial conditions; therefore, one initial guess could be to start with the nominal values.

### 5.2 Detecting IDH

The method could overcome the current weakness of estimating IDH during UF based on changes in HCT or protein and provide an alternative way of estimating a drop in blood volume and subsequent risk of IDH. Altering UF based on blood volume change could lead to a decrease in IDH [5]. As shown in Fig 3 and 5 (F), the method provides a prediction for the decline, as a curve, in blood volume during UF, which cannot be provided in the current dialysis practices. Since high UF rates, which are related to IHD, result in a sharper decline in blood volume, lower UF rates and extended time by clinicians can lead to increase blood volume curves and prevent IDH.

With the constant UF profile, we observed a rapid reduction in the blood volume before reaching the dry weight (see Fig 3, F). This observation is commonly associated with the constant UF profile because it uses a high constant rate throughout the dialysis session. However, the blood volume under the stepwise UF profile was slightly increased at the end of the session due to the decrease in the UF rates (see Fig 5, F). This remark can be seen when a designed UF profile is applied, in which the UF rate is objectively decreased toward the end of the session to control IDH [10, 29].

Another advantage of this approach is monitoring the reduction in the extracellular fluid overload during UF to reach a target dry weight and avoid fluid depletion, ultimately preventing IDH. Fig. 3 and 5 (C, D) illustrate the decrease of intravascular (plasma) and interstitial fluid volumes due to UF. The reduction in plasma volume and interstitial fluid volume under the constant UF profile reflects a significant drop in blood volume and the vascular refilling rate to compensate for the removed fluid volume during the treatment. We will consider these results in future work.

### 5.3 Limitations and Future Directions

The limitation of this study includes that this work presents the first attempt to apply this new approach to clinical data used in dialysis practices. We anticipate that the distribution of the estimated parameters will only be clear from larger dialysis data sets. But, here we have applied this approach for two data sets, including constant and stepwise UF profiles.

Future work will be focusing on clinical studies. We will apply this method to large dialysis data sets to obtain useful information for designing advanced treatments.

The proposed method could assist in diagnosing IDH events during the UF process. Using this new method in dialysis practices, we could predict the change in both blood and fluid volumes ahead of time. Our plan is to integrate this method into the existing dialysis machine to predict IDH events, which has not been done yet.

Finally, coupling this method with feedback-control systems to simultaneously estimate model parameters, update fluid and blood volumes, and predict IDH evens during dialysis treatment, is another future goal. In this feedback framework, blood volume is not only influenced by UF but also by the patient’s position or by taking a meal or drink during dialysis. We will implement external variables into the model to account for these circumstances.

## 6 Conclusions

This study presents a new approach for estimating blood and fluid volumes and vascular refilling-related parameters from dialysis data. The method predicts the changes in these volumes during UF. Unlike the other estimation methods, our method is not reliant on restricted UF or dilution protocols and can be used with different data lengths. This approach combines a nonlinear fluid volume model, an error quantization technique, and the UKF algorithm.

We validate the estimation result in four forms. First, we validate the result by implementing the Fisher information matrix to quantify error estimation. We showed that the blood and fluid volumes and the capillary filtration coefficient can be accurately estimated from a short data segment. However, since the hydrostatic capillary pressure parameters are insensitive to HCT measurements, the estimated uncertainty for these parameters was wide. Second, we use a short data segment to estimate the parameters and test the result by matching the measured HCT, in the rest of the data, using the modeled HCT. A good agreement between the measured HCT and the modeled HCT was obtained. Third, we showed that the estimation result using different data lengths are numerically very close, and the difference between the model’s responses is negligible. Last, we validate the estimation result by implementing the estimated parameters into the model and using it with the UKF algorithm to predict the HCT measurements during UF. We showed that a precise prediction of these measurements can be observed when these parameters are estimated correctly. This result is confirmed for both constant and stepwise UF profiles. Additionally, we analyzed the sensitivity of the estimation result to initial conditions. We found that the estimation result does not depend highly on initial conditions and is closely centered around the highest disturbed values. Finally, we provided a physiological understanding that our method can be used to detect IDH progression by interoperating the change in the blood and fluid volumes and assessing the vascular refilling rate during UF.

The direct impact of this study is the development of a new approach for estimating blood volume and fluid overload before dialysis and predicting the change in these variables during the UF process. This method is ready to implement into dialysis machines providing a better understanding of fluid volume management and monitoring patients in ESKD.

## Declarations

Conflict-of-interest: The authors have no conflicts of interest and agree with the manuscript’s contents.

Funding: The authors have no relevant financial or non-financial interests to disclose.

## Acknowledgements

The authors would like to thank Prof. Yossi Chait, University of Massachusetts Amherst, for his help and support.

## Authors’ Biographies

**Rammah Abohtyra**

**Rammah Abohtyra** received his M.S. and Ph.D. degrees in electrical engineering from Colorado School of Mines Golden, CO, USA, in 2013 and 2015, respectively. He is currently a Postdoc Scholar at the Pennsylvania State University, University Park, PA, USA. His primary research focuses on leveraging biological modeling, control system, and system identification to improve health problems, including fluid volume management in kidney failure, glycemic management, and neurobiological diseases.

**Tyrone Vincent**

**Tyrone Vincent** received his M.S. and Ph.D. degrees in electrical engineering from the University of Michigan, Ann Arbor, MI, USA, in 1994 and 1997, respectively. He is currently a Professor in the Department of Electrical Engineering and Computer Science at the Colorado School of Mines, Golden, CO, USA. His research interests include system identification, estimation, and control with applications in materials processing, energy systems, and biomedical engineering.

## Appendix A Fluid Volume Model

The equations for *P*_*c*_ (hydrostatic capillary pressure), *P*_*i*_ (interstitial pressure), *π*_*p*_ (plasma colloid osmotic pressure), *π*_*i*_ (interstitial colloid osmotic pressure) in (3b) are given by

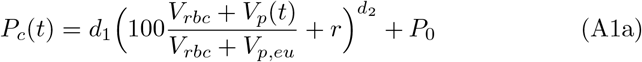

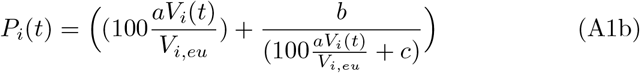

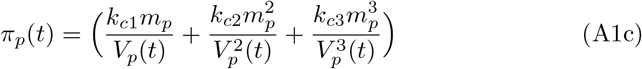

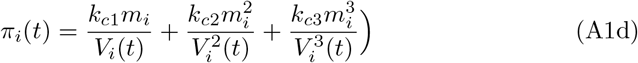

where *V*_*p,eu*_, *V*_*i,eu*_ denote euhydration (normal) plasma and interstitial volume for 70 kg patient, respectively; *P*_0_ is an offset pressure; *m*_*p*_ is plasma protein mass; *m*_*i*_ is interstitial protein mass, all are listed in Table A1, in which *a, b, c, d*_1_, *d*_2_, *r, m*_*p*_, *m*_*i*_, *k*_*c*1_, *k*_*c*2_ and *k*_*c*3_ are nominal parameters of model (1).

**Table A1.** The nominal parameters of the nonlinear two-compartment model.

| Parameter | Value | Parameter | Value |
| --- | --- | --- | --- |
| $a$ | 0.006 | $K_f$ L/(min*mmHg) | 0.0057 |
| $b$ | -198 | $m_p$ (g) | 210 |
| $c$ | -45 | $m_i$ (g) | 210 |
| $d_1$ | 0.01 | $P_o$ (mmHg) | 13.128 |
| $r$ | -30 | $V_{i,eu}$ (L) | 11 |
| $d_2$ | 1.46 | $V_{p,eu}$ (L) | 3 |
| $g$ | 0.045 | $k_{c1}$ | 0.21 |
| $h$ | 0.7672 | $k_{c2}$ | 0.0016 |
| $l$ | 0.045 | $k_{c3}$ | 9e-6 |
| $V_{rbc}(L)$ | 2 | - | - |

## Notes

### Competing Interest Statement

The authors have declared no competing interest.

### Summary of Updates

Updated version 1-20-2023

## References

[1] US Department of Health and Human Services, Centers for Disease Control and Prevention: Chronic Kidney Disease in the United States. [Online; accessed 19-August-2022] (2021). https://www.niddk.nih.gov/health-information/health-statistics/kidney-disease

[2] Krenn, S., Schmiedecker, M., Schneditz, D., Hödlmoser, S., Mayer, C.C., Wassertheurer, S., Omic, H., Schernhammer, E., Wabel, P., Hecking, M.: Feasibility of dialysate bolus-based absolute blood volume estimation in maintenance hemodialysis patients. Frontiers in medicine 9 (2022)

[3] Dasgupta, I., Farrington, K., Davies, S.J., Davenport, A., Mitra, S.: Uk national survey of practice patterns of fluid volume management in haemodialysis patients: a need for evidence. Blood purification 41(4), 324–331 (2016)

[4] Kanbay, M., Ertuglu, L.A., Afsar, B., Ozdogan, E., Siriopol, D., Covic, A., Basile, C., Ortiz, A.: An update review of intradialytic hypotension: concept, risk factors, clinical implications and management. Clinical Kidney Journal 13(6), 981–993 (2020)

[5] Sars, B., van der Sande, F.M., Kooman, J.P.: Intradialytic hypotension: mechanisms and outcome. Blood purification 49(1-2), 158–167 (2020)

[6] Palmer, B.F., Henrich, W.L.: Recent advances in the prevention and management of intradialytic hypotension. Journal of the American Society of Nephrology 19(1), 8–11 (2008)

[7] Daugirdas, J.T.: Pathophysiology of dialysis hypotension: an update. American journal of kidney diseases 38(4), 11–17 (2001)

[8] Thijssen, S., Kappel, F., Kotanko, P.: Absolute blood volume in hemodialysis patients: why is it relevant, and how to measure it. Blood purification 35(1-3), 63–71 (2013)

[9] Samandari, H., Schneditz, D., Germain, M.J., Horowitz, J., Hollot, C.V., Chait, Y.: A variable-volume kinetic model to estimate absolute blood volume in dialysis patients using dialysate dilution protocol. ASAIO Journal (American Society for Artificial Internal Organs: 1992) 64(1), 77 (2018)

[10] Abohtyra, R., Hollot, C., Germain, M.G., Chait, Y., Horowitz, J.: Personalized ultrafiltration profiles to minimize intradialytic hypotension in end-stage renal disease. In: 2018 IEEE Conference on Decision and Control (CDC), pp. 309–314 (2018). IEEE

[11] Pstras, L., Waniewski, J., Wojcik-Zaluska, A., Zaluska, W.: Relative blood volume changes during haemodialysis estimated from haemoconcentration markers. Scientific Reports 10(1), 1–9 (2020)

[12] Kron, S., Schneditz, D., Keane, D.F., Leimbach, T., Kron, J.: An improved method to estimate absolute blood volume based on dialysate dilution. Artificial Organs 45(9), 359–363 (2021)

[13] Kron, J., Schneditz, D., Leimbach, T., Aign, S., Kron, S.: A simple and feasible method to determine absolute blood volume in hemodialysis patients in clinical practice. Blood purification 38(3-4), 180–187 (2014)

[14] Schneditz, D., Schilcher, G., Ribitsch, W., Krisper, P., Haditsch, B., Kron, J.: On-line dialysate infusion to estimate absolute blood volume in dialysis patients. ASAIO Journal 60(4), 436–442 (2014)

[15] Kuhlmann, M.K., Zhu, F., Seibert, E., Levin, N.W.: Bioimpedance, dry weight and blood pressure control: new methods and consequences. Current opinion in nephrology and hypertension 14(6), 543–549 (2005)

[16] Agarwal, R., Weir, M.R.: Dry-weight: a concept revisited in an effort to avoid medication-directed approaches for blood pressure control in hemodialysis patients. Clinical Journal of the American Society of Nephrology 5(7), 1255–1260 (2010)

[17] Canaud, B., Chazot, C., Koomans, J., Collins, A.: Fluid and hemodynamic management in hemodialysis patients: challenges and opportunities. Brazilian Journal of Nephrology 41, 550–559 (2019)

[18] Silva, A.M., Heymsfield, S.B., Gallagher, D., Albu, J., Pi-Sunyer, X.F., Pierson Jr, R.N., Wang, J., Heshka, S., Sardinha, L.B., Wang, Z.: Evaluation of between-methods agreement of extracellular water measurements in adults and children. The American journal of clinical nutrition 88(2), 315–323 (2008)

[19] Zhu, F., Levin, N.W.: Dry weight and measurements methods. In: Penido, M.G. (ed.) Hemodialysis. IntechOpen, Rijeka (2011). Chap. 16. https://doi.org/10.5772/23868. https://doi.org/10.5772/23868

[20] Abbas, S.R., Zhu, F., Levin, N.W.: Bioimpedance can solve problems of fluid overload. Journal of Renal Nutrition 25(2), 234–237 (2015)

[21] Piccoli, A., et al.: Identification of operational clues to dry weight prescription in hemodialysis using bioimpedance vector analysis. Kidney international 53(4), 1036–1043 (1998)

[22] Guyton, A.C., Hall, J.: The body fluid compartments: extracellular and intracellular fluids; interstitial fluid and edema. Textbook of medical physiology 11, 293 (2000)

[23] Wiig, H., Swartz, M.A.: Interstitial fluid and lymph formation and transport: physiological regulation and roles in inflammation and cancer. Physiological reviews 92(3), 1005–1060 (2012)

[24] Kron, S., Schneditz, D., Leimbach, T., Aign, S., Kron, J.: Vascular refilling is independent of volume overload in hemodialysis with moderate ultrafiltration requirements. Hemodialysis International 20(3), 484–491 (2016)

[25] Schneditz, D., Roob, J., Oswald, M., Pogglitsch, H., Moser, M., Kenner, T., Binswanger, U.: Nature and rate of vascular refilling during hemodialysis and ultrafiltration. Kidney international 42(6), 1425–1433 (1992)

[26] Lindsay, R.M., Shulman, T., Prakash, S., Nesrallah, G., Kiaii, M.: Hemodynamic and volume changes during hemodialysis. Hemodialysis International 7(3), 204–208 (2003)

[27] Zhu, F., Schneditz, D., Wang, E., Martin, K., Morris, A., Levin, N.: Validation of changes in extracellular volume measured during hemodialysis using a segmental bioimpedance technique. ASAIO journal (American Society for Artificial Internal Organs: 1992) 44(5), 541–5 (1998)

[28] Rodriguez, H.J., Domenici, R., Diroll, A., Goykhman, I.: Assessment of dry weight by monitoring changes in blood volume during hemodialysis using crit-line. Kidney international 68(2), 854–861 (2005)

[29] Abohtyra, R., Chait, Y., Germain, M.J., Hollot, C.V., Horowitz, J.: Individualization of ultrafiltration in hemodialysis. IEEE Transactions on Biomedical Engineering 66(8), 2174–2181 (2019). https://doi.org/10.1109/TBME.2018.2884931

[30] Abohtyra, R.M., Hollot, C., Horowitz, J., Germain, M., Chait, Y.: Designing robust ultrafiltration rate profiles based on identifying fluid volume model parameters during hemodialysis. In: Dynamic Systems and Control Conference, vol. 58271, pp. 001–08006 (2017). American Society of Mechanical Engineers

[31] Fleming, S., Wilkinson, J., Aldridge, C., Greenwood, R., Muggleston, S., Baker, L., Cattell, W.: Dialysis-induced change in erythrocyte volume: effect on change in blood volume calculated from packed cell volume. Clinical nephrology 29(2), 63–68 (1988)

[32] Hamidi, M., Roshangar, F., Khosroshahi, H.T., Hassankhani, H., Ghafourifard, M., Sarbakhsh, P., et al.: Comparison of the effect of linear and step-wise sodium and ultrafiltration profiling on dialysis adequacy in patients undergoing hemodialysis. Saudi Journal of Kidney Diseases and Transplantation 31(1), 44 (2020)

[33] Flythe, J.E., Tugman, M.J., Narendra, J.H., Assimon, M.M., Li, Q., Wang, Y., Brunelli, S.M., Hinderliter, A.L.: Effect of ultrafiltration profiling on outcomes among maintenance hemodialysis patients: a pilot randomized crossover trial. Journal of Nephrology 34(1), 113–123 (2021)

[34] Abohtyra, R.M., Chait, Y.: New algorithm to design real time optimal and robust ultrafiltration rates in chronic kidney disease to prevent cardiovascular morbidity and mortality. In: Dynamic Systems and Control Conference, vol. 51890, pp. 001–11005 (2018). American Society of Mechanical Engineers

[35] Levenberg, K.: A method for the solution of certain non-linear problems in least squares. Quarterly of applied mathematics 2(2), 164–168 (1944)

[36] Julier, S.J., Uhlmann, J.K.: Unscented filtering and nonlinear estimation. Proceedings of the IEEE 92(3), 401–422 (2004)

[37] Van Trees, H.L.: Detection, Estimation, and Modulation Theory, Part I: Detection, Estimation, and Linear Modulation Theory. John Wiley & Sons, Hoboken, NJ (2004)

[38] Refaai, M.A., Goldstein, J.N., Lee, M.L., Durn, B.L., Milling Jr, T.J., Sarode, R.: Increased risk of volume overload with plasma compared with four-factor prothrombin complex concentrate for urgent vitamin k antagonist reversal. Transfusion 55(11), 2722–2729 (2015)

[39] Renkin, E.M.: Some consequences of capillary permeability to macromolecules: Starling’s hypothesis reconsidered. American Journal of Physiology-Heart and Circulatory Physiology 250(5), 706–710 (1986)

[40] He, P., Hu, J., Huang, C.: P1162 red blood cell distribution width and peritoneal dialysis-associated peritonitis prognosis. Nephrology Dialysis Transplantation 35(Supplement 3), 142–1162 (2020)

[41] Marano, S., Marano, M., Pecchia, L.: Frontiers in hemodialysis part II: Toward personalized and optimized therapy. Biomedical Signal Processing and Control 61, 102029 (2020)

[42] Marano, S., Marano, M., Gennari, F.J.: A new approach to bicarbonate addition during hemodialysis: Testing model predictions in a patient cohort. IEEE Access 10, 17473–17483 (2022)

